# Development of a simple method for differential delivery of volatile anaesthetics to the spinal cord of the rabbit

**DOI:** 10.1101/785253

**Authors:** Peng Zhang, Yao Li, Ting Xu

## Abstract

Emulsified volatile anaesthetics can be easily injected into the circulation and eliminated from blood to lungs. Taking advantage of this unique pharmacokinetics, we aimed to develop a less trauma model for preferential delivery of volatile anaesthetics to the spinal cord with an intact CNS and circulatory system in the rabbit. Sixteen New Zealand White rabbits were randomly assigned to the isoflurane and sevoflurane group. A catheter for delivering emulsified isoflurane (8mg/kg/h) or sevoflurane (12mg/kg/h) to the spinal cord was placed into the descending aorta in all rabbits. The concentration and partial pressure of isoflurane and sevoflurane in the jugular and femoral vein were measured. Our results showed that the partial pressure of isoflurane was 3.91±1.11 mmHg in the jugular vein and 12.61±1.60 mmHg (1.0MAC) in the femoral vein. The partial pressure of sevoflurane was 3.89±1.00 mmHg in the jugular vein and 19.92±1.84mmHg (1.0MAC) in the femoral vein. There was significant difference regarding the partial pressure of isoflurane and sevoflurane between jugular and femoral veins in two groups (both *P*<0.001). In conclusion, a simple method has been successfully developed to selectively deliver isoflurane and sevoflurane to the spinal cord in the rabbit. Before acting on the brain, 69% isoflurane and 81% sevoflurane were removed from body via lungs. This method can be used to investigate the pharmacological properties of volatile anesthetics.

## Induction

Volatile anaesthetics can induce a variety of reversible, clinically important effects such as amnesia, hypnosis and immobility[1, 2]. To determine if an effect is due to brain, spinal cord, peripheral nerves or both, we need a differential anaesthetic delivery model[3, 4]. To solve this problem, previous researches have introduced several selectively anaesthetized models in the goat, dog and rabbit, using cardiopulmonary bypass technology [5–7] But, these models limited by the severe trauma which could destroy the physiological structure of the cerebral and spinal cord circulation of experimental animals.

Rabbits have a unique spinal cord circulation in that each spinal cord segment is supplied by a single corresponding radicular artery arising from aorta[8, 9].In addition, the blood supply to the thoracolumbar region (below T3) of spinal cord originate from the thoracic and abdominal aorta in the rabbit [10–12]. Emulsified volatile anaesthetic is that liquid anaesthetic dissolves in an emulsion and produces anaesthesia as effectively as liquid one [13, 14]. What’s important is that, emulsified volatile anaesthetic can be directly injected into the circulation as same as intravenous drugs and eliminated via the lungs[15, 16]. Based on these findings, we aimed to establish a less trauma model for differential delivery of volatile anaesthetics (isoflurane and sevoflurane) to the spinal cord with an intact CNS and circulatory system in the rabbit.

## Materials and methods

### Emulsified isoflurane and sevoflurane preparation

Isoflurane and sevoflurane emulsion were prepared according to a previously described formula[17, 18]. In brief, 1.6mL of liquid isoflurane or sevoflurane (Abbott, Shanghai, China) and 18.4mL of 30% intralipid (Baxter, Suzhou, China) were injected into a sealed sterile vial. Then the vial was violently shaken for 10min by a vibrator to solubilise isoflurane or sevoflurane into the intralipid. The 8% emulsified isoflurane and sevoflurane were stored in 4°C refrigerator before use.

### Animal preparation and surgical procedures

The study was approved by the Institutional Animal Care and Use Committee of Sichuan Provincial People’s Hospital. Sixteen New Zealand White rabbits (male and female) weighting 2.0-3.0kg were randomly assigned to the isoflurane and sevoflurane group. After intravenous injection of 30 mg/kg pentobarbital sodium into the left marginal ear vein, a ID 3.0# cuffed endotracheal tube was inserted. All rabbits were mechanical ventilated with 95% O_2_ using an animal ventilator (Chengdu Techman Software Co.Ltd, Chengdu, China), with the tidal volume 8ml/kg, 40 breaths/min, to maintain the arterial carbon dioxide pressure (PaCO2) between 35 to 45 mmHg. A 22-gauge IV catheter (BD company, Sandy Utach, USA) was individually inserted into the left central ear and femoral artery to monitor the blood pressure in the upper and lower torso of the rabbit in each group.

An epidural catheter (TuoRen Medical Instrument Co., Ltd, Xinxiang, China) which was used to deliver emulsified isoflurane or sevoflurane to the spinal cord was placed into the descending aorta from the right femoral artery (Fig 1). The change of resistance was monitored during the catheter passing cranially along the aorta. The catheter would be withdrawn 0.5-1.0 cm, when the increased resistance was recorded. The place of catheter tip was confirmed by autopsy after the experiment. In our study, the rabbit would be excluded from data analysis if the tip was located in the aortic arch level. The length of catheter between catheter tip and inguinal fold was measured.

**Fig 1.**
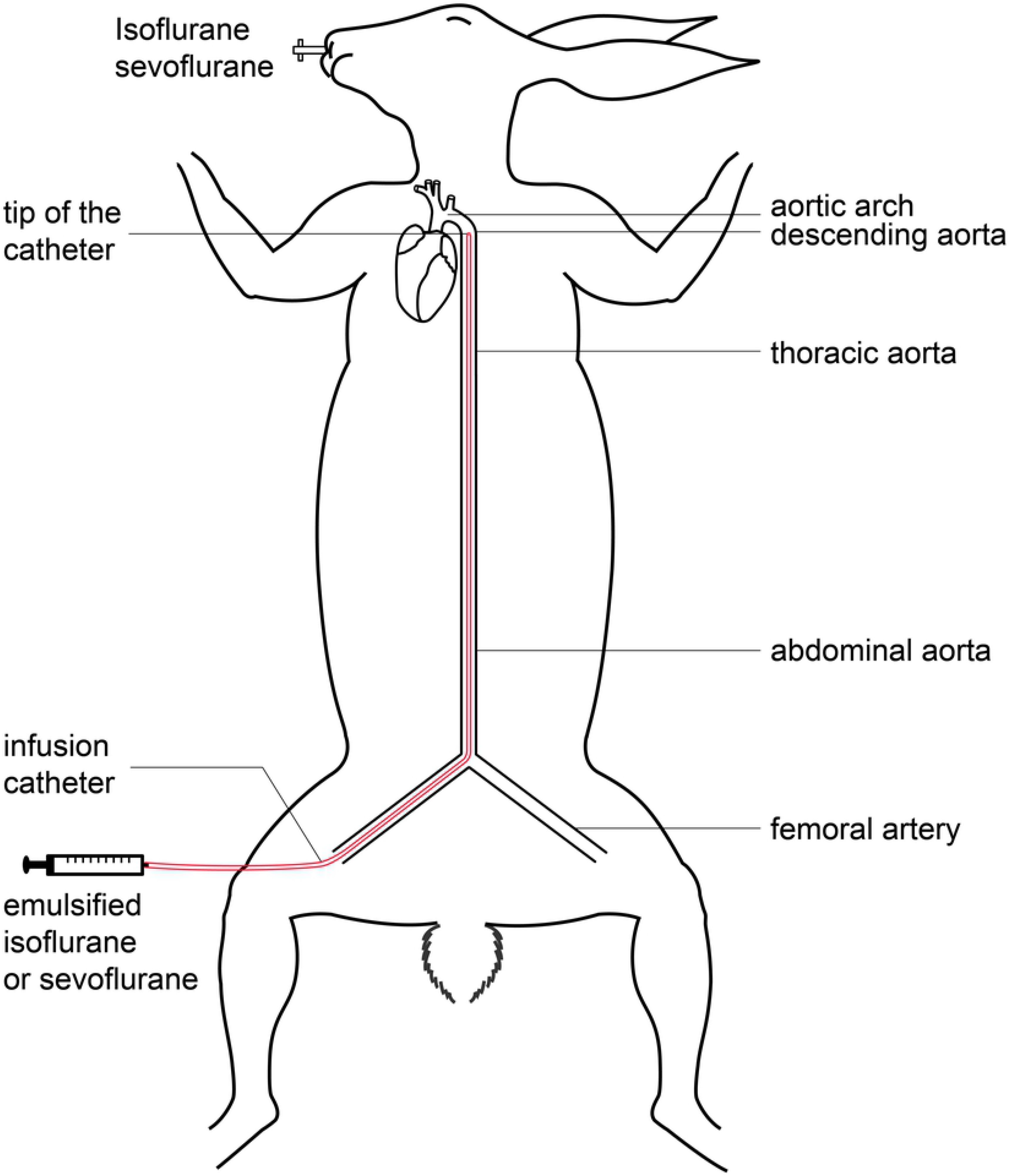
Diagram of the differential anesthetic delivery rabbit model

The infusion rate was 8mg/kg/h in the isoflurane group and 12mg/kg/h in the sevoflurane group by a microinfusion pump. When the end-tidal isoflurane concentration (E_T_ISO) or end-tidal sevoflurane concentration (E_T_SEVO) was stable and maintained for 20min, emulsified volatile anaesthetic was considered to reach steady state condition[15]. There were two methods to avoid re-breaching of isoflurane or sevoflurane into the brain: (a.) a higher inspiratory flow 2L/min (normal minute ventilation (MV) of rabbit is about 1L/min) (b.) an anaesthetic absorber made of activated charcoal in the inspiratory limb of the circuit.

Lactated Ringer’s solution (6ml/kg/h) was administered through left marginal ear vein in all rabbits. Rectum temperature was monitored and maintained at 37.0±1.0°C. During the experiment, heart rate (HR), mean arterial pressure (MAP), electrocardiograph (ECG) and pulse oxygen saturation (SpO_2_) were measured with the BL-420E+ Data Acquisition and Analysis System (Chengdu Techman Software Co.Ltd, Chengdu, China). The E_T_ISO, E_T_SEVO and E_T_CO_2_ (end-tidal CO_2_) were monitored with an anesthetic gas analyzer M1026B (Philips Medizin Systeme Boblingen GmbH, Boblingen, Germany).

### Collection of blood samples and gas chromatograph

Once the end-tidal concentration of isoflurane or sevoflurane was stable and maintained for 20min, 5-6ml blood was rapidly withdrawn from the jugular and femoral vein to determine the isoflurane or sevoflurane concentration (C) in the blood, using a gas chromatograph Agilent 4890D (Tegent Technology Ltd., Hong Kong, China) with the two-stage headspace equilibration methods[15]. The partial pressure of isoflurane and sevoflurane (P) in the different blood samples were calculated by the equation of *P*= (C/λ_b/g_×760 mmHg) (λ_b/g_ =blood/gas partition coefficient). The C and P of all samples was measured by a technician who was blinded to the samples. In our previous study, the partial pressure of 1.0 MAC (minimum alveolar concentration) isoflurane was 11.66±1.10 mmHg and the partial pressure of 1.0 MAC sevoflurane was 18.86±1.12 mmHg.

### Statistical analysis

Statistical analyses were performed using SPSS 22.0. Each continuous variable was analyzed for its normal distribution and reported as mean ± SD. Repeated measures ANOVA was used to evaluate MAP and HR among different time points in two groups. Pearson correlation coefficient was used to measure the linear relationship between λ_b/g_ and the volume (mL) of the emulsified isoflurane or sevoflurane consumed. Student’s t test and χ^2^ test were also measured in our study. A *P* value of less than 0.05 was considered statistically significant. A Bonferroni correction was made when necessary to correct for multiple testing.

## Results

Sixteen rabbits finished the experiment and all data were analyzed. The mean weights of the rabbits were 2.38±0.26 kg in the isoflurane group and 2.46±0.24 kg in the sevoflurane group. No significant difference was found between two groups (*P*=0.532). The mean lengths of the inserted catheter (between catheter tip and inguinal fold) were 25.13±1.07 cm in the isoflurane group and 25.36±0.86 cm in the sevoflurane group. In the isoflurane group, two catheters (25%) were located at T_2_ and six (75%) at T_3_. In the sevoflurane group, four catheters (50%) were placed at T_2_ and four (50%) at T_3_ (Table 1). There was no significant difference with regard to the length of the catheter (*P*=0.608) and position of the catheter tips (*P*=0.631) between two groups.

**Table 1.** The position of catheter tips in two groups

| Group | T<sub>2</sub> | T<sub>3</sub> | Total |
| --- | --- | --- | --- |
| Isoflurane | 2 (25%) | 6 (75%) | 8 |
| Sevoflurane | 4 (50%) | 4 (50%) | 8 |
| Total | 6 | 10 | 16 |
| *P* value | 0.608 |  |  |

No significant difference was found regarding MAP of the central ear artery among three different time points in two groups (*P*=0.169 for isoflurane group; *P*=0.390 for sevoflurane group). But MAP of the femoral artery at the midpoint of emulsified anaesthetics infusion was significantly reduced compared with other two time points (after induction and end of the study) in two groups (both *P*<0.001). The trend of MAP change between central ear artery and femoral artery was significantly different during the whole experiment in two groups (both *P*<0.001). However, there was no significant difference in the MAP between central ear artery and femoral artery during the whole experiment in two groups (*P*=0.062 for isoflurane group; *P*=0.169 for sevoflurane group) (Table 2). No significant difference was found with regard to HR among three time points in the isoflurane group (*P*=0.091) and sevoflurane group (*P*=0.076) (Table 3).

**Table 2.** The mean atrial pressure (MAP) of femoral and central ear artery among different time points in two groups (mean ± SD)

| Group | Position | After induction | Midpoint of infusion | End of the study |
| --- | --- | --- | --- | --- |
| Isoflurane | Central ear artery | 90.4±2.3 | 86.3±5.5 | 87.5±4.5 |
| | Femoral artery | 92.3±3.6 | 74.6±6.1<sup>#&</sup> | 88.4±4.4 |
| Sevoflurane | Central ear artery | 88.1±5.1 | 84.6±7.5 | 84.3±5.4 |
| | Femoral artery | 90.3±5.0 | 71.4±4.3<sup>#&</sup> | 86.1±4.8 |

<sup>#</sup> $P < 0.001$ , midpoint of infusion vs after induction
<sup>&</sup> $P < 0.001$ , midpoint of infusion vs end of the study

**Table 3.** The heart rate (HR) among different time points in two groups (mean ± SD)

| Group | After induction | Midpoint of infusion | End of the study |
| --- | --- | --- | --- |
| Isoflurane | 203±11 | 201±5 | 212±11 |
| Sevoflurane | 215±11 | 207±9 | 220±13 |

The partial pressure of isoflurane was 3.91±1.11 mmHg in the jugular vein and 12.61±1.60 mmHg (1.0MAC) in the femoral vein. The partial pressure of sevoflurane was 3.89±1.00 mmHg in the jugular vein and 19.92±1.84mmHg (1.0MAC) in the femoral vein. There was significant difference regarding the partial pressure of isoflurane and sevoflurane between jugular and femoral vein in two groups (both *P*<0.001). The ratio of partial pressure of isoflurane and sevoflurane between jugular vein and femoral vein was showed in the Table 4.

**Table 4.** The partial pressure of isoflurane and sevoflurane (mmHg) in the jugular and femoral vein in two groups (mean ± SD)

| Group | P<sub>j</sub><br>jugular vein | P<sub>f</sub><br>femoral vein | Ratio1<br>(P<sub>j</sub>/ P<sub>f</sub>)% | Ratio2<br>(P<sub>f</sub>/ P<sub>j</sub>) | *P* value |
| --- | --- | --- | --- | --- | --- |
| Emulsified isoflurane<br>(8mg/kg/h) | 3.91±1.11 | 12.61±1.60 | 31.28±9.17% | 3.48±1.13 | <0.001 |
| Emulsified sevoflurane<br>(12mg/kg/h) | 3.89±1.00 | 19.92±1.84 | 19.40±4.46% | 5.34±1.18 | <0.001 |

There was a significant positive correlation between λ_b/g_ and volume of the emulsified anaesthetics delivered (r=0.935, *P*<0.001 for isoflurane group; r=0.919, *P*=0.001 for sevoflurane group). The Linear regression equation was y=0.0869x+0.9688 (R^2^=0.8723) in the isoflurane group (Fig 2) and y=0.0526x+0.3889 (R^2^=0.8452) in the sevoflurane group (Fig 3) (y=volume and x= λ_b/g_).

**Fig 2.**
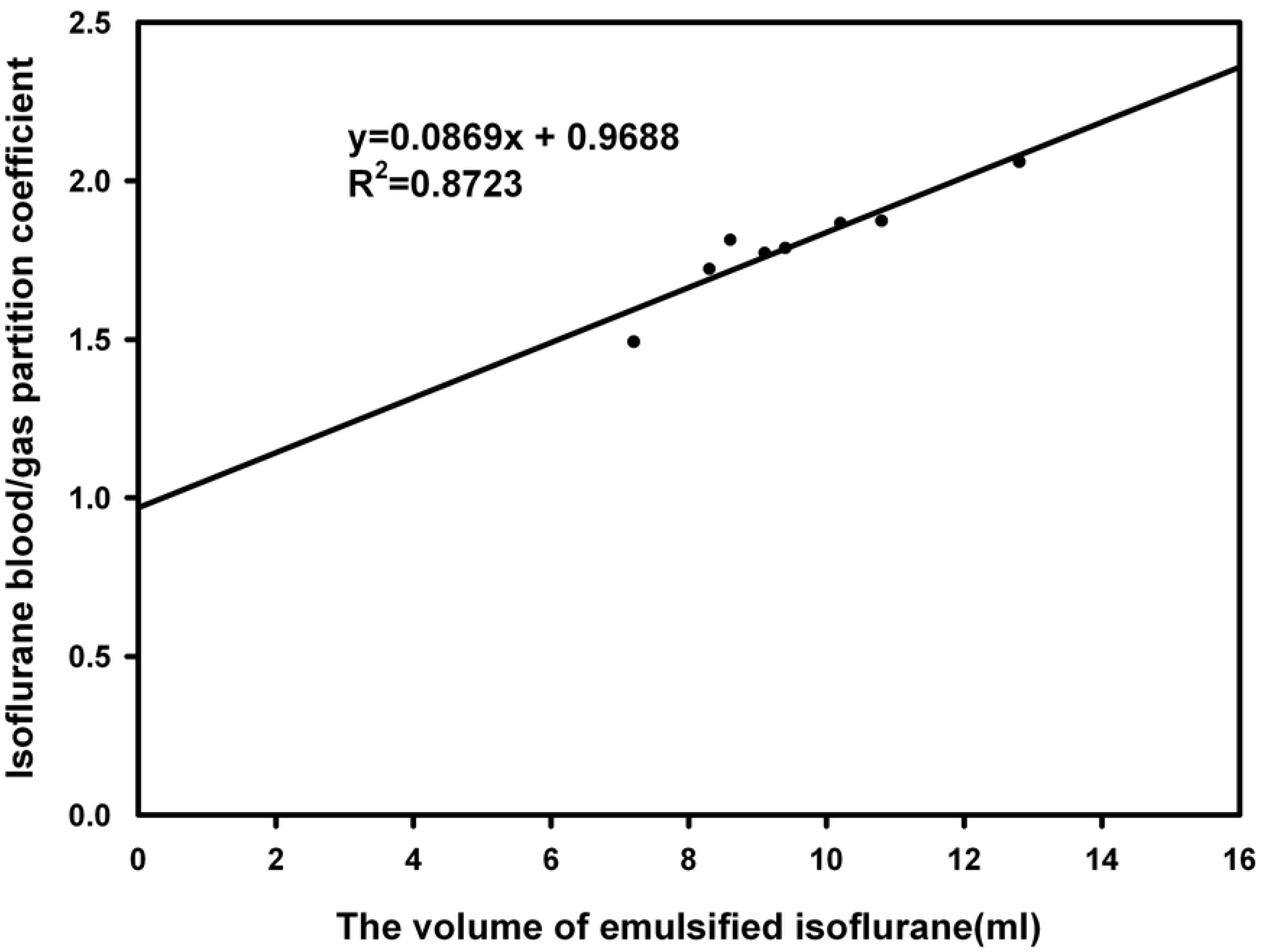
The positive correlation between isoflurane blood/gas partition coefficient and the volume of emulsified isoflurane delivered

**Fig 3.**
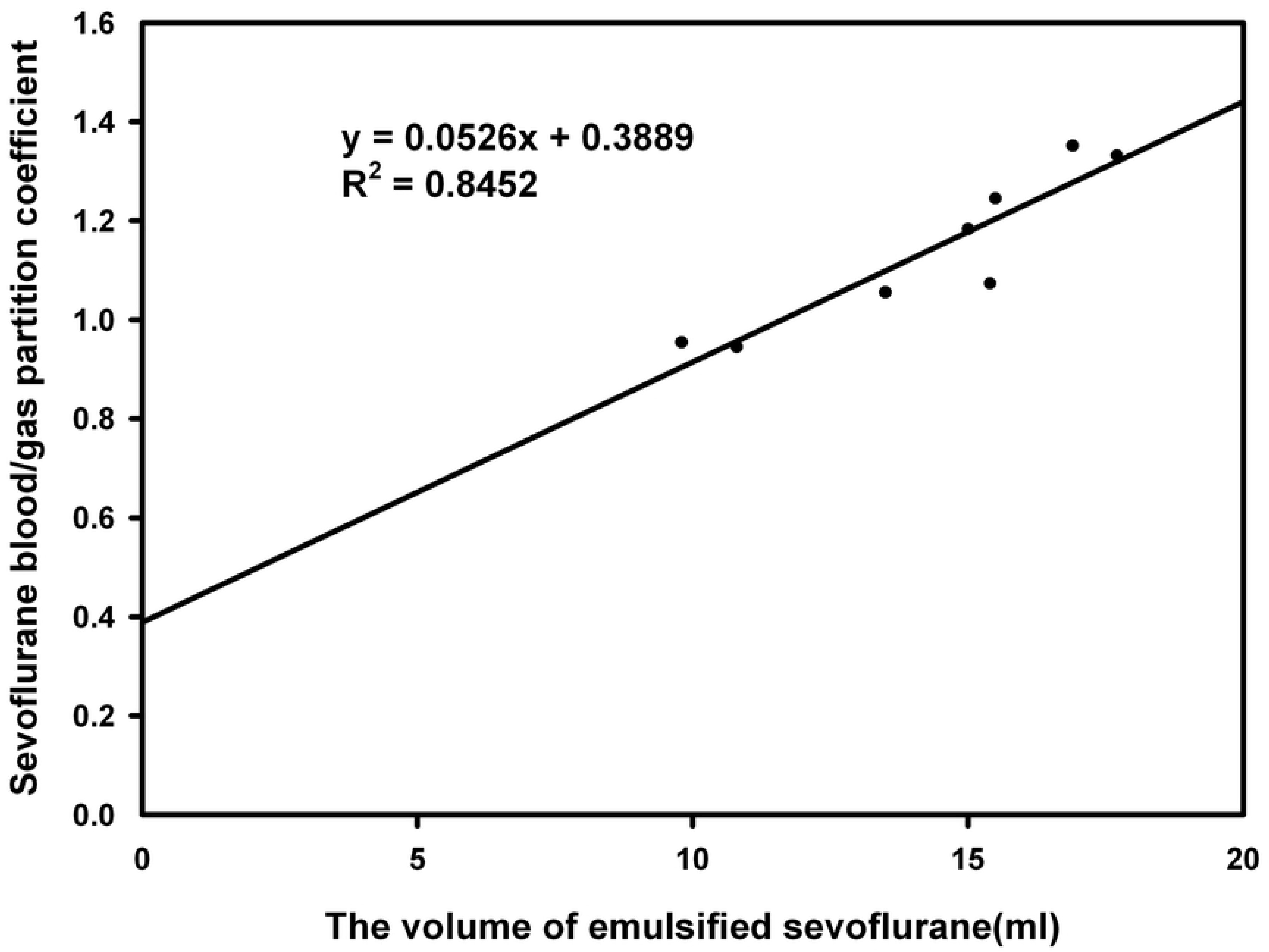
The positive correlation between sevoflurane blood/gas partition coefficient and the volume of emulsified sevoflurane delivered

## Discussion

The rabbit, with its medium size and homosegmental blood supply to the spinal cord, make it an ideal animal for selectively anaesthetized models [10–12]. Due to the catheter inserted into the descending aorta, isoflurane and sevoflurane emulsion were able to be preferentially delivered to the thoracolumbar (below T_4_) region of the spinal cord. But before acting on the brain, the majority of it were prior eliminated via the lungs. According to our results, the rabbit model could exhale 69% isoflurane or 81% sevoflurane by lungs, before delivering to the brain. Therefore, the lower torso circulation of the rabbit was successfully bypassed from the upper torso part of the body in our study.

When volatile anaesthetic reach steady state conditions in the body, the partial pressure in the alveolar, blood and central nervous system is presumed to be in equilibrium with each parts [15, 19]. In our study, to be specific, the partial pressure in the jugular and femoral vein can reflect that in the brain and spinal cord tissue respectively under steady state condition. Our results showed that the isoflurane partial pressure in the femoral vein was about 3.5-fold more than that in the jugular vein, and the ratio of sevoflurane was 5.3-fold. In other words, the partial pressure of isoflurane and sevoflurane in the spinal cord were 3.5-fold and 5.3-fold more than that in the brain tissue. Thus, our model could differentially deliver emulsified isoflurane and sevoflurane to the spinal cord in the rabbit.

In *Yang’s* goat model, using similar technologies, emulsified isoflurane was selectively delivered to the spinal cord, with an approximate 46% reduction in the isoflurane partial pressure in the brain [18]. However, the partial pressure of isoflurane in the brain was only about 31% of that in the spinal cord, while the ratio of sevoflurane was 19% in our study. Our rabbit model seemed more effective in removing anaesthetics than previous goat model. There were several possible reasons: (a.) The oxygen consumption per kilogram of rabbit is quite larger than that of goat, and MV/kg of rabbit is about 2 times than that of goat [20]. It has established that increasing MV might contribute to remove volatile anaesthetics from blood to lungs [21]. (b.)Two methods mentioned (higher inspiratory flow and anaesthetic absorber) were used to avoid re-breaching of isoflurane or sevoflurane (c.) In our study, sevoflurane was easier to be eliminated from body than isoflurane, because of lower blood gas solubility (λ_b/g_ of sevoflurane=0.069; λ_b/g_ of isoflurane=1.37) [22].

It is well known that λ_b/g_ of 30% intralipid is remarkably larger than that of isoflurane or sevoflurane. So λ_b/g_ of isoflurane and sevoflurane would significantly increase with infusion of lipid emulsion in our study. Our results indicated that 1 ml of 8% emulsified isoflurane increased λ_b/g_ by 0.0869 at 37°C, and 0.0526 for sevoflurane λ_b/g_. Some discrepancy researches showed that 1 ml of 8% emulsified isoflurane raised λ_b/g_ by 0.0176 in the goat [18] and 0.0139 in the dog[15]. Obviously, the blood volume of rabbits is significantly less than that of goats or dogs[21]. Based on “volume fraction partition coefficient” theory, when 1ml of lipid emulsion was administrated to those three animals, the change of rabbit λb/g would be probably the biggest one [23]. But there was no similar report published regarding λ_b/g_ of sevoflurane.

In our study, because of blood supply limitations to the spinal cored, only thoracolumbar region (below T_3_), not the entire spinal cord, could be selectively anaesthetized during aortic delivery of emulsified isoflurane and sevoflurane. Though rabbit brain could not completely be separated from the entire spinal cord, our research still had few several advantages. Firstly, because of less trauma, compared with cardiopulmonary bypass, the integrity of circulation and CNS (central nervous system) can remain intact in our study[6, 7, 24]. Secondly, rabbits are common experimental animals and may have financial benefit. At the end of the study, all rabbits awoke and no fatalities was found. This confirmed that infusion of emulsified isoflurane or sevoflurane produced a safe and reversible anaesthetic effect with no apparent pathological damage to the CNS and circulatory system.

In conclusion, a simple method has been successfully established to differential delivery of isoflurane and sevoflurane to the spinal cord with an intact CNS and circulatory system in rabbits. This model permitted to eliminate 69% isoflurane and 81% sevoflurane from blood to lungs, before acting on the brain. This allows us to determine if an anaesthesia action is due to brain or spinal cord effects. In addition, rely on rabbit unique homosegmental blood supply to the spinal cord, this model can also be used to preferential delivery of anaesthetics to the lumbar or sacro-coccygeal spinal cord, thereby permitting to investigate the effects of different each spinal cord segment on anaesthesia action.

## Author Contributions

Animal experiment: Peng Zhang and Yao Li

Data analysis: Yao Li

Writing-original draft: Peng Zhang

Writing-review and editing: Ting Xu

